# Identification of fetal liver stroma in spectral cytometry using the parameter autofluorescence

**DOI:** 10.1101/2021.09.29.462345

**Authors:** Marcia Mesquita Peixoto, Francisca Soares-da-Silva, Sandrine Schmutz, Marie-Pierre Mailhe, Sophie Novault, Ana Cumano, Cedric Ait-Mansour

## Abstract

The fetal liver is the main hematopoietic organ during embryonic development. The fetal liver is also the unique anatomical site where hematopoietic stem cells expand before colonizing the bone marrow, where they ensure life-long blood cell production and become mostly resting. The identification of the different cell types that comprise the hematopoietic stroma in the fetal liver is essential to understand the signals required for the expansion and differentiation of the hematopoietic stem cells. We used a panel of monoclonal antibodies to identify fetal liver stromal cells in a 5-laser equipped spectral flow cytometry analyzer. The “Autofluorescence Finder” of SONY ID7000 software identified two distinct autofluorescence emission spectra. Using autofluorescence as a fluorescence parameter we could assign the two autofluorescent signals to three distinct cell types and identified surface markers that characterize these populations. We found that one autofluorescent population corresponds to hepatoblasts and cholangiocytes whereas the other expresses mesenchymal transcripts and was identified as stellate cells. Importantly, after birth, autofluorescence becomes the unique identifying property of hepatoblasts because mature cholangiocytes are no longer autofluorescent.

These results show that autofluorescence used as a parameter in spectral flow cytometry is a useful tool to identify new cell subsets that are difficult to analyze in conventional flow cytometry.

## Introduction

Flow cytometry (FCM) has rapidly evolved over the past few decades (1). The development of more advanced technologies including mass cytometry (2), imaging flow cytometry (3), genomic flow cytometry (4) and spectral flow cytometry (5) has expanded our ability to study immune responses, by characterizing more cellular parameters simultaneously in single cells with higher resolution than ever before. FCM characterizes physical and fluorescent properties of cells in suspension by using fluorochrome-conjugated antibodies to measure proteins expressed by distinct immune cell subpopulations (6). Spectral flow cytometry was first proposed by J. Paul Robinson at Purdue University in 2004. The first commercial instrument was launched by Sony Biotechnology in 2012, using prisms along with PMT detectors to collect and amplify light beyond the capability of conventional flow cytometers. The Sony SP 6800 (Sony Inc) was previously described (7), it is equipped with three lasers 405/488/638nm and two pinholes such that two excitation lasers can be used in a given experiment. Using the 405/488nm excitation lasers we assembled a 19-florescent probe panel to analyze immune cells in the mouse adult spleen (8,9). Moreover, analysis of antibody-stained preparations of embryonic, neonatal and adult heart, which is a highly auto-fluorescent tissue, allowed the robust identification of new surface markers of different cardiac populations (10). Recently, the Sony ID 7000 became available with different laser configurations that can be remodeled. The LE-ID7000E used in this study has 5 lasers 355/405/488/561/637nm, PMT can be independently regulated and there are 7 pinholes allowing for additional lasers to be installed. The ID7000 Software analysis allows the easy detection of auto-fluorescence (AF) that can be incorporated in the panel as an independent fluorescent parameter.

The fetal liver (FL) is a highly auto-fluorescent tissue where hepatic and hematopoietic cells differentiate alongside and where hepatic and mesenchymal cells play, yet ill-defined, roles in hematopoietic differentiation (11) and in the establishment of the hematopoietic stem cell compartment (12). In addition, although surface markers for different populations have been identified, a global analysis of the FL stroma using epithelial and mesenchymal markers has not been undertaken.

Here we sought to analyze the non-hematopoietic stromal compartment in mouse FL using a 14-fluorochrome antibody panel and distinct AF parameters that were unique in the identification of cell populations. We found two major AF signals at embryonic day (E)18.5 of mouse gestation. One AF signal has a high intensity in the 355nm laser and is also detected at lower levels in the 405nm laser. It defines stellate cells that accumulate autofluorescent vitamin A in cytoplasmic granules. The second AF signal is visible in the 355nm laser, in the 405nm and also with a lower intensity in the 488 and 561nm lasers and marks hepatoblasts and cholangiocytes. Importantly, AF is an invaluable tool to distinguish hepatoblasts from hepatocytes because they lose distinctive markers but maintain AF, after birth. In addition, defining the true fluorescence associated with each autofluorescent population helped design a sorting strategy to be used in conventional FCM. These results indicate that AF treated as a fluorescence parameter in spectral flow cytometry is essential to the characterization of cell subsets in solid tissues.

## Materials and Methods

### Cell suspension

E18.5 fetal livers (FL) P3 and P9 livers were dissected under a binocular magnifying lens. Cells were recovered in Hanks’ balanced-salt solution (HBSS) supplemented with 1% fetal calf serum (FCS) (Gibco) or RPMI medium (Gibco) supplemented with 10% FCS (P3 and P9) and treated with 0.05 mg/ml Liberase TH (Roche) and 0.2 mg/ml DNAse (Sigma) for 6-8 minutes at 37°C. Cells were resuspended by gentle pipetting. Before staining, cell suspensions were filtered with a 100 μm cell strainer (BD).

### Flow cytometry and cell sorting

Liver cells were depleted of Ter119^+^CD45^+^ CD71^+^ CD117^+^ cells by magnetic cell separation using LS MACS Columns (Miltenyi Biotec). Cell suspensions were stained for 20-30 min at 4ºC in the dark with antibodies listed in Table 1. Biotinylated antibodies were detected by incubation for 15 min at 4°C in the dark with streptavidin. Stained cells were analyzed on Sony ID7000 and were sorted with a BD FACSAria III (BD Biosciences) according to the guidelines for the use of FCM and cell sorting (6). Before the analysis the Sony ID7000 was calibrated using the daily QC and the performance 8-peak beads and the FACSAria III was calibrated with CST beads following the manufacturer’s protocols. Data were analyzed with the ID7000 (Sony, Inc) or FlowJo (v.10.7.3, BD Biosciences) software.

**Table 1.** List of the antibodies used in this study.

Fetal liver cells analyzed at E12.5, E14.5 and E18.5
|  |  |  |
| --- | --- | --- |
| SAV | PE-Cy5 | CD45, TER119, CD71, CD117 (all hematopoietic cells) biotin |
| Dead | PI |  |
| **CD324** | **PE-Cy7** | (Epithelial-Cadherin) hepatoblasts, hepatocytes and cholangiocytes |
| CD31 | PerCP-Cy5.5 | (Platelet and endothelial cell adhesion molecule) endothelial cells |
| **CD166** | **FITC** | (Activated Leukocyte Cell Adhesion Molecule) hepatoblasts and stellate cells |
| **CD140a** | **APC** | (Platelet derived growth factor receptor $\alpha$) stellate cells and pericytes |
| Gp38 | APC-Cy7 | (Podoplanin) mesenchymal cells |
| **NG2** | **PE** | (chondroitin sulphate proteoglycan 4) pericytes |
| Sca-1 | BV711 | (Stem Cell antigen 1) endothelial cells |
| CD146 | BV510 | (melanoma cell adhesion molecule) endothelial cells and mesenchymal cells |
| CD51 | BV421 | (Integrin $\alpha$ V) mesenchymal cells |
| **CD326** | **BV605** | (Epithelial cell adhesion molecule) cholangiocytes (bile duct cells) |
| CD54 | BUV395 | (Intercellular Adhesion Molecule 1) hepatoblasts and endothelial cells |
| Thy1.2 | AF700 | (cluster of differentiation 90) mesenchymal cells |

### Gene expression by RT-PCR

Cells were sorted directly into lysis buffer and mRNA was extracted with an RNeasy Micro Kit (Qiagen). After extraction mRNA was reverse-transcribed into cDNA with PrimeScript RT Reagent Kit (Takara Bio), followed by quantitative PCR with Power SYBR Green PCR Master Mix (Applied Biosystems). Primers used were as follows: Actb FW: GCTTCTTTGCAGCTCCTTCGT; Actb RV: ATCGTCATCCATGGCGAACT; Afp FW: CTTCCCTCATCCTCCTGCTAC; Afp RV: ACAAACTGGGTAAAGGTGATGG; Alb FW: TGCTTTTTCCAGGGGTGTGTT; Alb RV: TTACTTCCTGCACTAATTTGGCA; Des FW: GTGGATGCAGCCACTCTAGC; Des RV: GTGGATGCAGCCACTCTAGC; Onecut1 FW: GCCCTGGAGCAAACTCAAGT; Onecut1 RV: TTGGACGGACGCTTATTTTCC; Pdgfrb FW: TTCCAGGAGTGATACCAGCTT; Pdgfrb RV: AGGGGGCGTGATGACTAGG; Sox9 FW: CTCCTCCACGAAGGGTCTCT; Sox 9 RV: AGGAAGCTGGCAGACCAGTA. qPCR reactions were performed on a Quantstudio3 thermocycler (Applied Biosystems), gene expression was normalized to that of β-actin and relative expression was calculated using the 2^-ΔCt^ method

### Quantification and Statistical Analysis

All results are shown as mean ± standard deviation (SD).

## Results

### A strategy to characterize non-hematopoietic cells populations in fetal liver

To characterize non-hematopoietic (hereafter referred to as stromal) populations in the FL we assembled a panel of 14 fluorescent dyes (Table 1) that included propidium iodide (dead cell exclusion) and 13 fluorescent proteins that mark cells belonging to the hepatic, endothelial and mesenchymal lineages (pericytes and liver fibroblasts also called stellate cells). The analysis was done in FL cell suspensions depleted of hematopoietic cells in a dump channel that includes the erythroid marker TER119, the pan-hematopoietic marker CD45, and CD71 and CD117 that together mark embryonic erythroid progenitors. Cells were depleted of hematopoietic cells, that represent more than 80% of total FL cells, by magnetic sorting. We analyzed FL cells isolated from embryonic day E 18.5 (one day prior to birth), and from 3 and 9 days post-birth (P). Figure 1A shows the spectra of emission of the cells after the elimination of hematopoietic and dead cells. Figure 1B shows the gating strategy used to identify known cell subsets with CD31 and CD146 defining endothelial cells, and, in non-endothelial cells, CD140a (PDGFra), Gp38 (podoplanin), and CD166 (ALCAM) define stellate cells as CD140a^+^Gp38^-^CD116^+^. In the Gp38^-^CD140a^-^ subset, CD326 and CD324 (E-Cadherin) (13) define cholangiocytes (CD324^+^CD326^+^)(14), hepatoblasts (CD324^high^CD326^-^) and hepatocytes (CD324^low^CD326^-^). This analysis shows that at E18.5 all major FL populations are detected, and hepatocytes are difficult to discriminate from the decreasing numbers of hepatoblasts that are undergoing differentiation.

**Figure 1.**
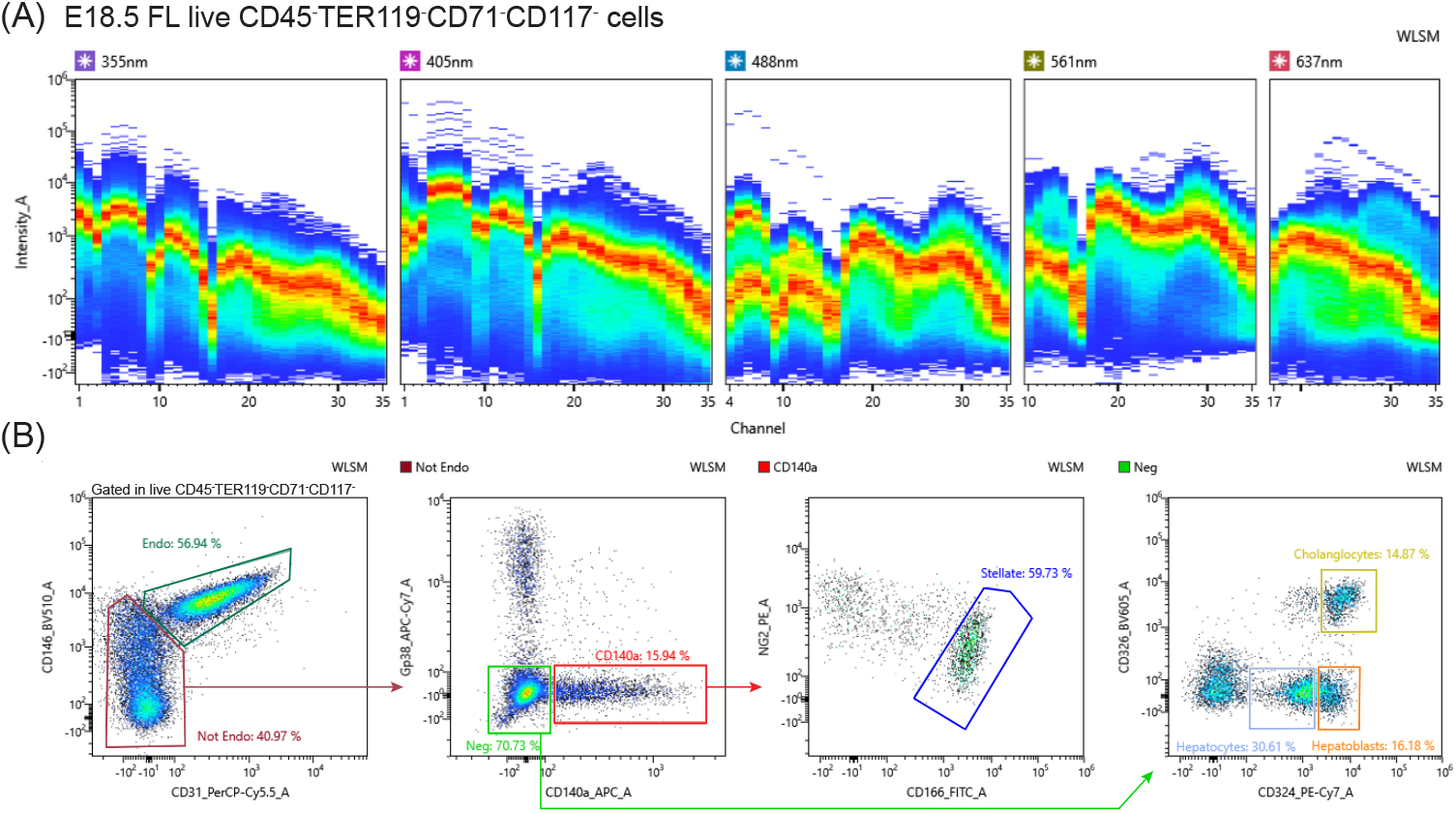
Gating strategy to identify stellate cells, hepatoblasts, hepatocytes and cholangiocytes. E18.5 mouse fetal liver cell suspensions were stained with biotin labelled antibodies recognizing CD45, TER119, CD71, CD117. Negative cells were magnetically sorted and labelled with the antibodies in Table 1. **A**. The spectra of emission of fetal liver cells after depletion (marked with streptavidin in a dump channel) and gated out of PI labelled cells. **B**. After unmixing, cells were manually gated to define endothelial and stellate cells. CD324 was defined as a marker of epithelial cells, cholangiocytes CD324^high^CD326^+^; hepatocytes CD324^low^CD326^-^; hepatoblasts CD324^high^CD326^-^.

### Two different autofluorescence patterns identified by the autofluorescence finder

AF is a property of many tissues defined as signal emission by cells excited at particular wavelengths, in the absence of specific fluorescence. AF can mask or distort true fluorescence and is usually subtracted to facilitate the fluorescence detection. The ID7000 spectral cytometer is equipped with an analytical software that provides a tool to identify independent AF signals called AF finder (Figure 2). In unstained cells this tool allowed the detection of two AF signals (Figure 2A). AF-1 is detected in the 355, and 405nm lasers with decreasing intensity and virtually undetected in the remaining lasers. AF-2 is detected with a high signal in the 355 and 405nm lasers, but also at a lower level in the 488 and 575 nm lasers and virtually undetected in the 461nm laser. The differences in the spectra of emission found in most laser channels suggested that these two AF signals correspond to two independent cell types.

**Figure 2.**
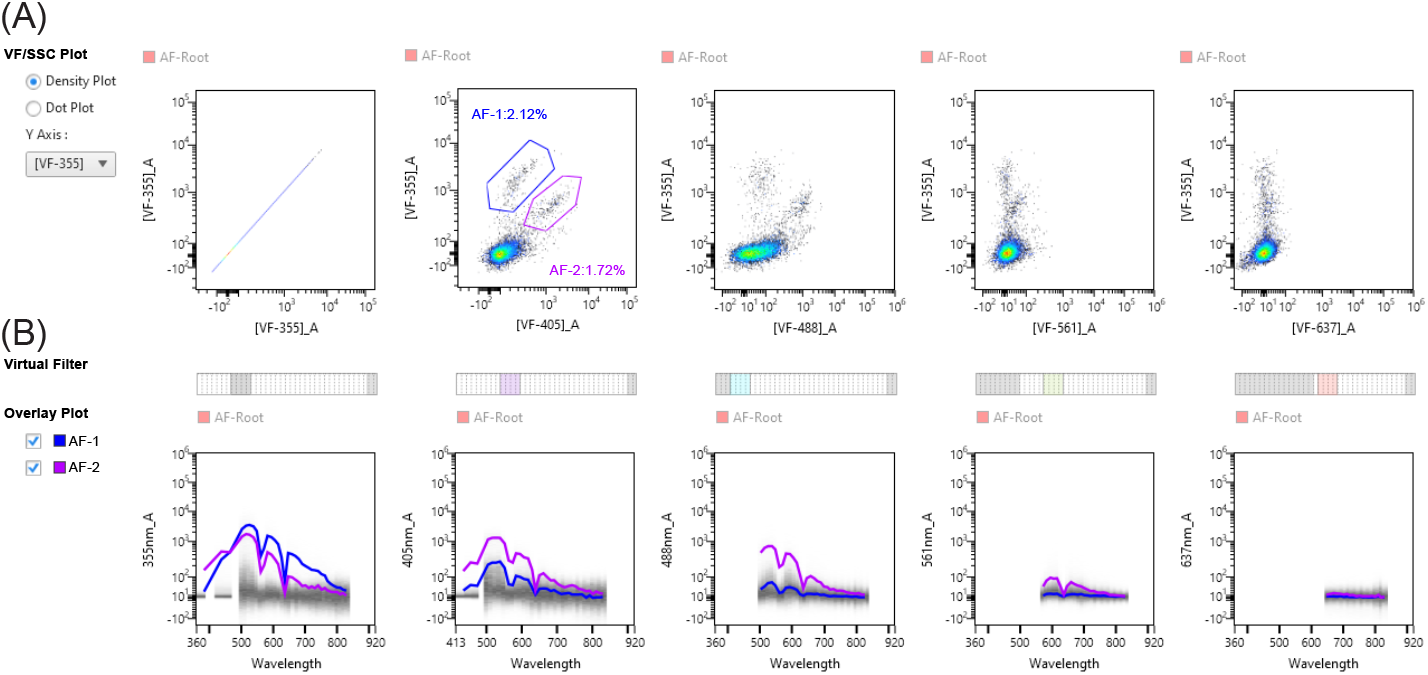
Autofluorescence finder. The analytical tool autofluorescence applied to unstained cells before unmixing. **A**. Autofluorescence is defined manually in cells that show fluorescence in the different lasers. **B**. The spectra of emission of the two defined autofluorescent (AF) signals in the different lasers.

After unmixing, we used AF-1 and AF-2 as independent parameters to define two different populations and proceeded to identify their identity (Figure 3 A-D). Figure 3 shows two independent gating strategies, one that analyzes cells within AF-1 (Figure 3B) compared to the conventional strategy previously shown in Figure 1 (Figure 3A). AF-1 cells correspond to a majority of CD31^-^ (85%), CD140a^+^ (85%) cells and CD116^+^ (95%) cells (Figure 3B) that define, with a high degree of enrichment, stellate cells.

**Figure 3.**
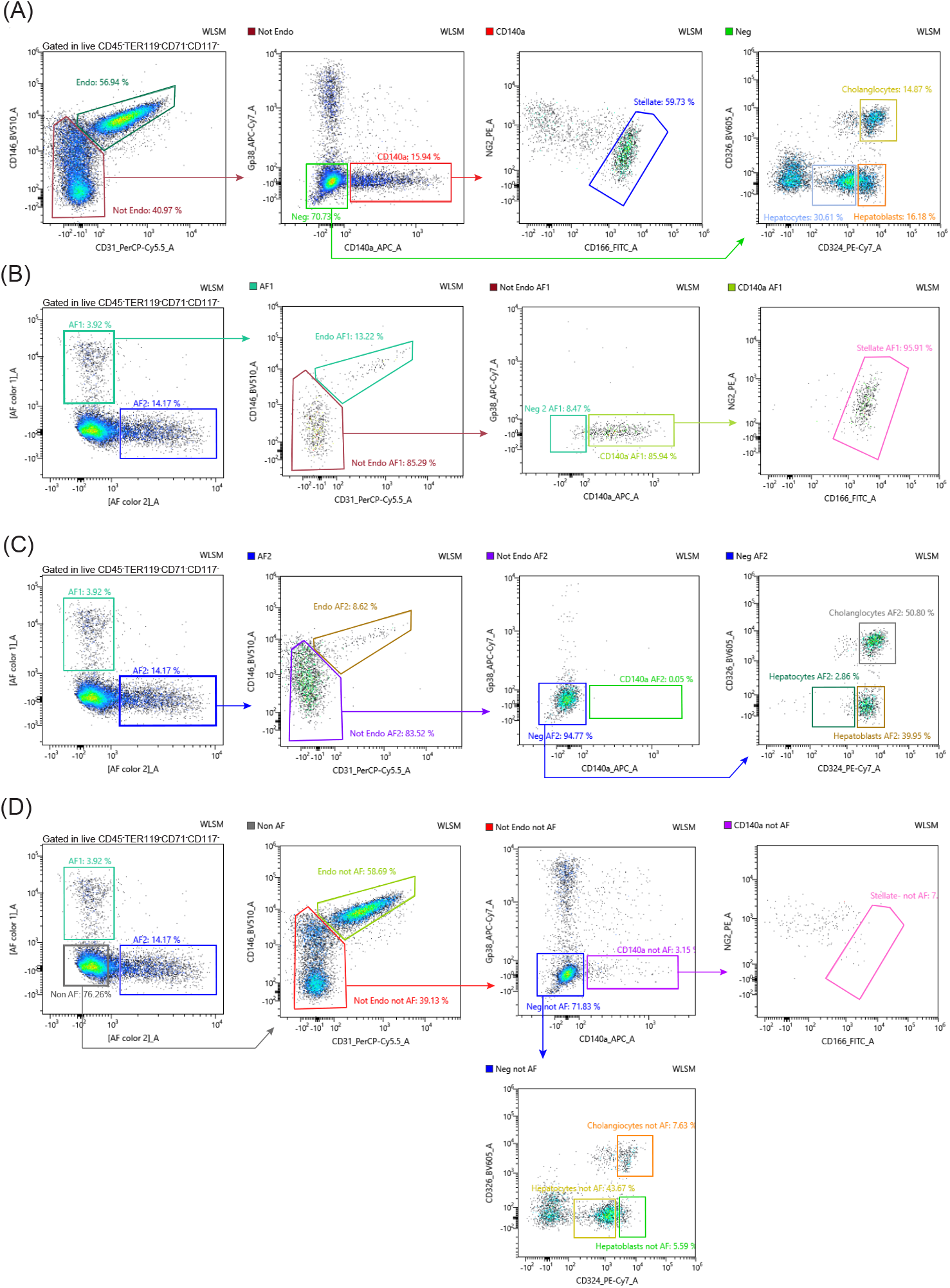
Two distinct autofluorescencent signals: AF-1 defines stellate cells and AF-2 defines a subset of hepatoblasts and cholangiocytes. After unmixing the two AF signals appear as two additional fluorescent parameters. **A**. E18.5 FL live CD45^-^TER119^-^CD71^-^CD117^-^ cells were analyzed as in Figure 1B. **B**. Analysis of E18.5 FL live CD45^-^TER119^-^CD71^-^CD117^-^AF1^+^ cells using the same strategy as in Figure 1B. **C**. Analysis of E18.5 FL live CD45^-^TER119^-^CD71^-^CD117^-^AF2^+^ cells using the same strategy as in Figure 1B. **D**. Analysis of E18.5 FL live CD45^-^TER119^-^CD71^-^CD117^-^AF1^-^AF2^-^ cells using the same strategy as in Figure 1B.

CD324^+^ hepatocytes and CD324^+^CD326^+^ cholangiocytes (bile duct cells) are known to derive during embryonic development from bipotent progenitors called hepatoblasts (15). AF-2 gated cells (Figure 3C) comprise a majority (83%) of CD31^-^ non-endothelial cells, 94% of CD140a^-^ gp38^-^ and >90% of CD324^high^ hepatoblasts and CD324^+^ CD326^+^ cholangiocytes (16). Interestingly, the analysis done in AF-2 did not detect the subset that expresses low levels of CD324 that we classify as hepatocytes (13)(Figure 3C). AF-2 is, therefore, an important parameter to distinguish hepatoblasts from hepatocytes, at this stage of development. When auto-fluorescent cells were gated out, stellate cells are virtually absent from the plots (Figure 3D) whereas a subset of hepatoblasts and cholangiocytes are still detected (30% and 50%, respectively) indicating that AF-2 distinguishes two different cell subsets (AF-2^+^ and AF-2^-^ cells) in each of these populations.

### Properties and identity of fetal liver stromal cells

To further characterize the AF of FL cells we determined whether this is a cell-intrinsic property or if it is lost upon manipulation we sorted hepatocytes, hepatoblasts and stellate cells from E18.5 FL using a simplified panel in a conventional flow cytometer (Figure 4A, presort). All populations before and after cell sorting were analyzed in spectral FCM (Figure 4B, sorted populations). Stellate cells (lower plots) show persistent levels of AF whereas hepatocytes (upper plots) remained negative for this parameter. As previously observed, hepatoblasts sorted as CD324^high^ cells yield a mixed population of AF-2^+^ and AF-2^-^ cell (middle plots). These results indicated that AF in fetal liver populations is a cell-specific property.

**Figure 4.**
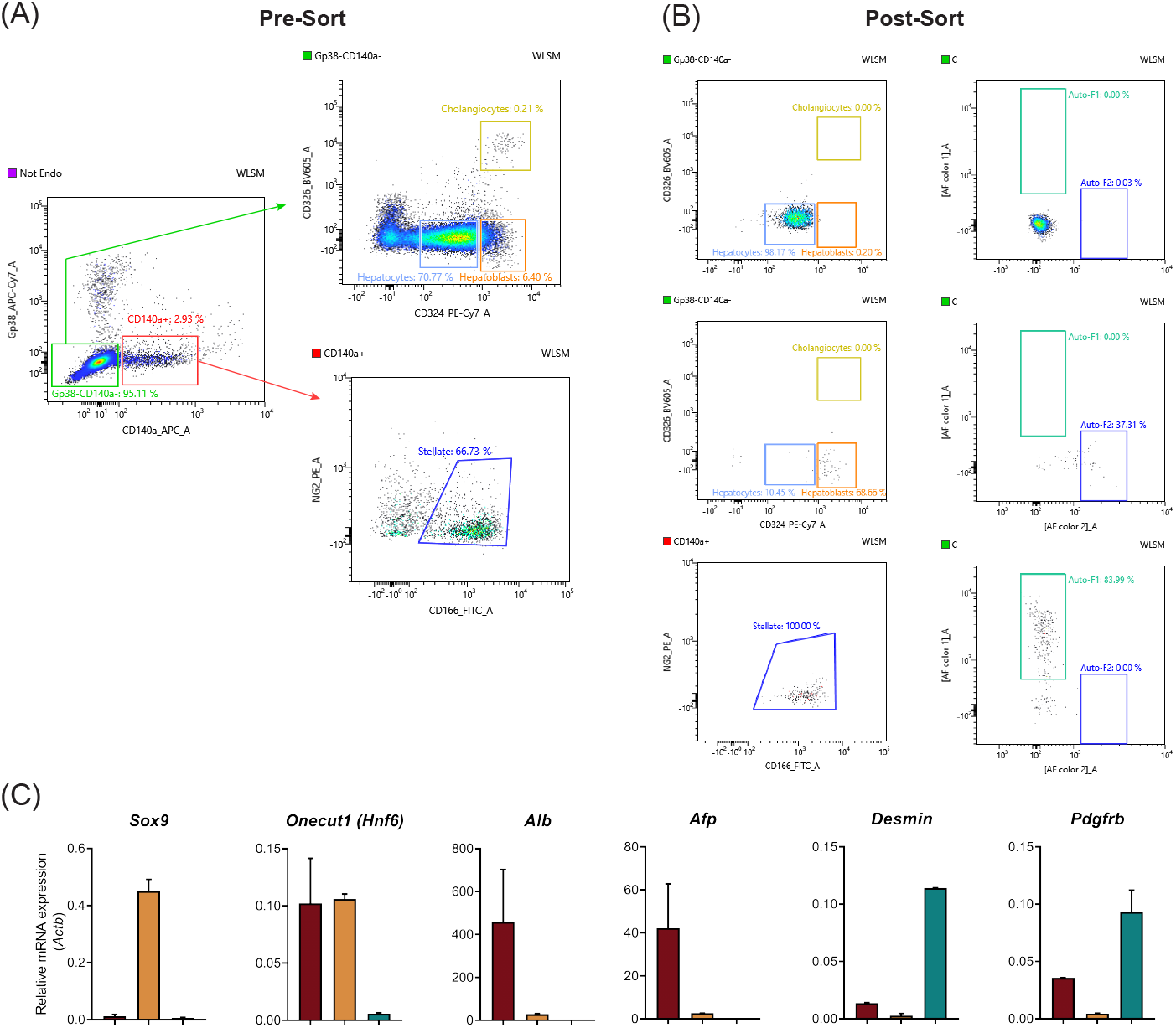
Autofluorescence is a cell intrinsic property. **A**. Fetal liver cells negative for the dump channel and for PI expressing cells were sorted after being stained with a simplified antibody panel including Gp38, CD140a, NG2, CD166, CD324 and CD326. Hepatocytes, hepatoblasts and stellate cells were sorted according to the strategy shown (pre-sort). Sorted cells were reanalyzed in the ID7000 spectral analyzer (sorted populations). Left plots show purity control, right plots show AF of sorted cells. **B**. Sorted cholangiocytes, hepatoblasts and stellate cells as in **A** were subjected quantitative RT-PCR in triplicate samples of two independent experiments for *Sox9*, *Onecut1*, *Alb*, *Afp*, *Desmin* and *Pdgfrb*, and normalized to the expression of β-actin used as house keeping gene. Data are represented as mean +/- SD

In other to unambiguously ascertain the lineage affiliation of these three cell subsets we subjected the cells sorted to quantitative RT-PCR. We show that cells classified as cholangiocytes by flow cytometry express the cholangiocytic markers *Sox9* and *Onecut 1* (*Hnf6*) (14), the hepatoblasts express *Onecut 1* (15), *Alb* and *Afp* whereas stellate cells express *Desmin* and *Pdgfrb* (Figure 4C) (17). This transcriptional profile is specific for the three subsets thus confirming the lineage affiliation inferred by the surface marker expression.

### Autofluorescence defines hepatoblasts after birth

Hepatoblasts decrease in numbers as development progresses, so we used AF-2 to detect these cells, after birth. However, hepatocytes that are the dominant liver population after birth are fragile epithelial cells, highly sensitive to manipulation *ex vivo* and to the disruption of the tight junctions, required for single-cell suspensions. Consequently, we detected decreasing numbers of hepatocytes with age. At P3 and P9 the levels of AF were very similar in spectrum and intensity to those seen at E18.5. At P3 and P9 AF-2 represents a majority of hepatoblasts with high expression of CD324, and by high AF-2 (Figure 5A). Interestingly, at these stages, most cholangiocytes lost expression of AF-2. This result is reinforced at P9 (Figure 5B) where gating in AF-2 yields 93% of hepatoblasts. These results further indicate that AF-2 is essential for the identification of hepatoblasts after birth.

**Figure 5.**
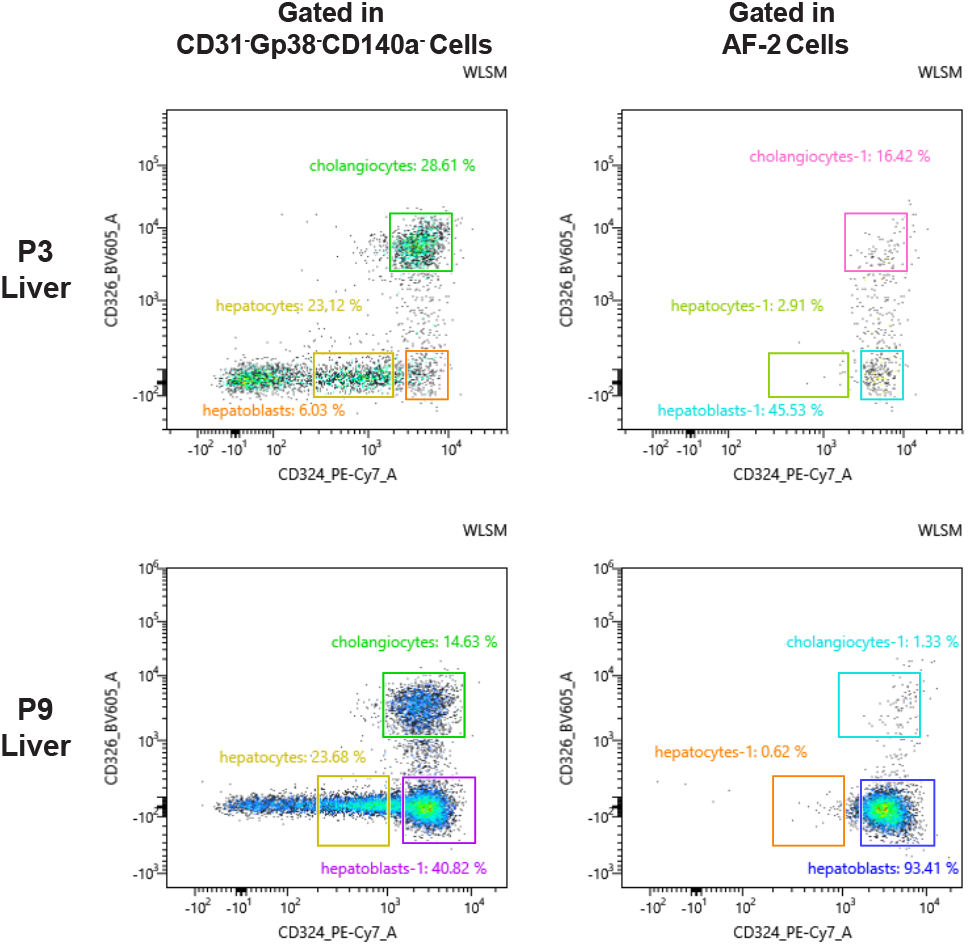
AF2 defines hepatoblasts after birth. Analysis of live CD45^-^TER119^-^CD71^-^CD117^-^ P3 and P9 liver cells in a plot of CD324/CD326 after the conventional gating strategy defined in Figure 1 (left plots) or on the cells gated for AF2 expression (right plots).

## Discussion

In this report we used a spectral flow cytometer equipped with 5 lasers to analyze a highly autofluorescent tissue, the FL, in single-cell suspensions. The software analysis of the Sony ID7000 is equipped with an analytical tool for easy detection of different autofluorescent signals. The unmixing algorithm subtracts the AF to find true fluorescent signals but importantly, by using the AF signals as independent parameters, it increased the panel of 14 fluorescent dyes into a 16-fluorescence panel.

The “Autofluorescence finder” tool identified independent AF signals that differed in shape and intensity in the different laser detectors. Gating on AF-1 we identified a dominant population with the phenotype that defines stellate cells, the fibroblasts specific to the liver. This population has previously been reported to be endowed with AF in a limited analysis using conventional FCM because stellate cells store vitamin A in cytoplasm granules (18). The second AF signal, AF-2, was found in epithelial cells (expressing CD324) that, at E18.5, could be subdivided into two subsets. The first expressed CD326 which is specific, at this stage, of newly differentiated cholangiocytes (16), and the second expressed the highest levels of CD324 while lacking CD326 and was consequently classified as hepatoblasts (13).

We found that the two independent AF signals in stromal FL populations were intrinsic properties of the cells which confers reliability to the analysis. The transcriptional profile of the AF populations unambiguously confirmed the lineage affiliation proposed by the surface marker analysis.

The AF-2 parameter allowed the distinction of hepatocytes from hepatoblasts (CD324^+^ CD326^-^ cells), by clearly demarcating the limits between the two cell types. Because hepatoblasts rapidly decrease after birth, as most proliferative activity in the adult liver is driven by hepatocytes, we analyzed liver cells after birth. We found that the two different AF signals persisted at these later time points with AF-2 marking a majority of hepatoblasts. Interestingly, most cholangiocytes lost AF-2 at P3 and virtually all were AF2^-^ at P9. AF-2 is therefore the only property specific for hepatoblasts, after birth. Although hepatocytes are the most numerous cells in the adult liver, they comprised less than 20% of the cell suspension analyzed here. This observation is consistent with the well-documented fragility of hepatocytes. The protocols currently recommended to isolate viable hepatocytes include enzymatic digestion similar to the procedure used in this study (19). Alternatively, a perfusion with an enzymatic solution is followed by a purification in a Percoll gradient (20). In any of these methods, 5-20 x10^6^ hepatocytes (19) (1-6%) can be obtained out of the 3×10^8^ hepatocytes in a mouse liver (21) indicating that no efficient procedure is currently available to isolate hepatocytes.

Our study presents several innovations: 1. The ID7000 software analysis provides an easy tool to detect independent AF signals; 2. The unmixing algorithm subtracts the AF signals to allow accurate true fluorescence detection, a feature already present in the SP6800; 3. Importantly, the analytical tool treats the AF signals as fluorescence and increases, therefore, the panel of analyzed parameters. We provide evidence that these advances will be invaluable to the characterization of single-cell suspensions obtained from of autofluorescent tissues, difficult to analyze in conventional FCM.

## Aknowledgements

We thank all members of the flow cytometry core facility, the members of the Institut Pasteur laboratory, Antonio Bandeira, Paulo Vieira, Rachel Golub and Pablo Pereira. This work was financed by the Institut Pasteur, INSERM, ANR (grant Twothyme and grant EPI-DEV), REVIVE Future Investment Program and Pasteur-Weizmann Foundation through grants to A.C. This work was financed by Portuguese funds through FCT/MCTES in the framework of the project PTDC/MED-OUT/32656/2017 (POCI-01-0145-FEDER-032656). F.S.S is funded by the REVIVE Future investments post-doctoral grant (Investissement d’Avenir; ANR-10-LABX-73). MP is funded by FCT grant SFRH/BD/143605/2019.

